# Mapping the Global Distribution of *Mus musculus*: Implications for Evolutionary Genetics

**DOI:** 10.1101/2024.07.09.602589

**Authors:** Alexis Garretson, Laura Blanco-Berdugo, Beth Dumont

## Abstract

House mice (*Mus musculus*) are a key biomedical research model and important vectors for disease transmission. In the wild, house mice are also an ecologically disruptive invasive species, and their activity is associated with significant economic and agricultural damage and cost. Despite the importance of house mice across these different contexts, the extent of their geographic distribution is not well understood. House mice are human commensals but are nonetheless sensitive to their prevailing environment, indicating that the range of human settlement cannot be used as a reliable proxy. Existing range maps for *Mus musculus* are based on minimum convex hulls informed by potentially biased sampling and do not 1) fully integrate large, digitized data documenting species occurrences, 2) provide insight into the likely species distribution in under-sampled regions, and 3) delineate internal structures of the range, including barriers to dispersal or unsuitable internal habitat. Consequently, we know little about the bioclimatic tolerance and environmental envelope occupied by this species. To address these unknowns, we leverage publicly available mouse sampling and biodiversity data to provide an updated range map of *Mus musculus* and define the environmental limits of the house mouse distribution. Using genetic data from public archives, we also model the genetic diversity of house mice across our newly updated range. Using these data, we visualize global genetic diversity trends and confirm the ancestral origins of *Mus musculus* to the region of the Indian subcontinent occupied by modern-day Pakistan and northwestern India. Taken together, our efforts highlight areas where house mice are predicted to be at their environmental tolerance limit, including regions where future sampling efforts may uncover mice with unique adaptive traits.

## Introduction

House mice (*Mus musculus*) are one of the most biomedically, economically, and ecologically important rodent species. House mice are the premier mammalian model system for biomedical research and an important zoonotic vector for diseases such as hantavirus, salmonellosis, leptospirosis, lymphocytic choriomeningitis (LCMV), tularemia, and plague. From an economic perspective, house mice cause wide-ranging impacts on food production through crop destruction and contamination of stored foods. Indeed, from 1930-2022, it is estimated that house mice caused nearly one billion dollars of property and crop damage [1]. Ecologically, house mice are an extremely successful invasive species [2–4]. They can compete with indigenous lizards, seabirds, invertebrates, and plant species [5–11], leading to displacement or perturbation of native species balance. Their significant impact across multiple domains underscores the importance of understanding global mouse distributions and habitat preferences.

*Mus musculus* (L., 1758) is thought to have originated in the northern Indian subcontinent [12,13]. The prevailing model of mouse demographic history holds that mice radiated from this ancestral region approximately 250,000-500,000 years ago, giving rise to the three core house mouse subspecies (Figure 1) [14,15]. The three subspecies are found in different geographic regions, with *M. m. domesticus* (Schwarz & Schwarz, 1943) in Western Europe and the Iranian Valley, *M. m. musculus* (L.) in Eastern Europe and Northern Asia, and *M. m. castaneus* (Waterhouse 1843) primarily in Central and Southeastern Asia [12,13,16]. Aided by historical and contemporary trading routes, house mice have expanded their range outside of these ancestral regions in recent history, colonizing both North and South America, Australasia, and numerous remote islands.

**Figure 1.**
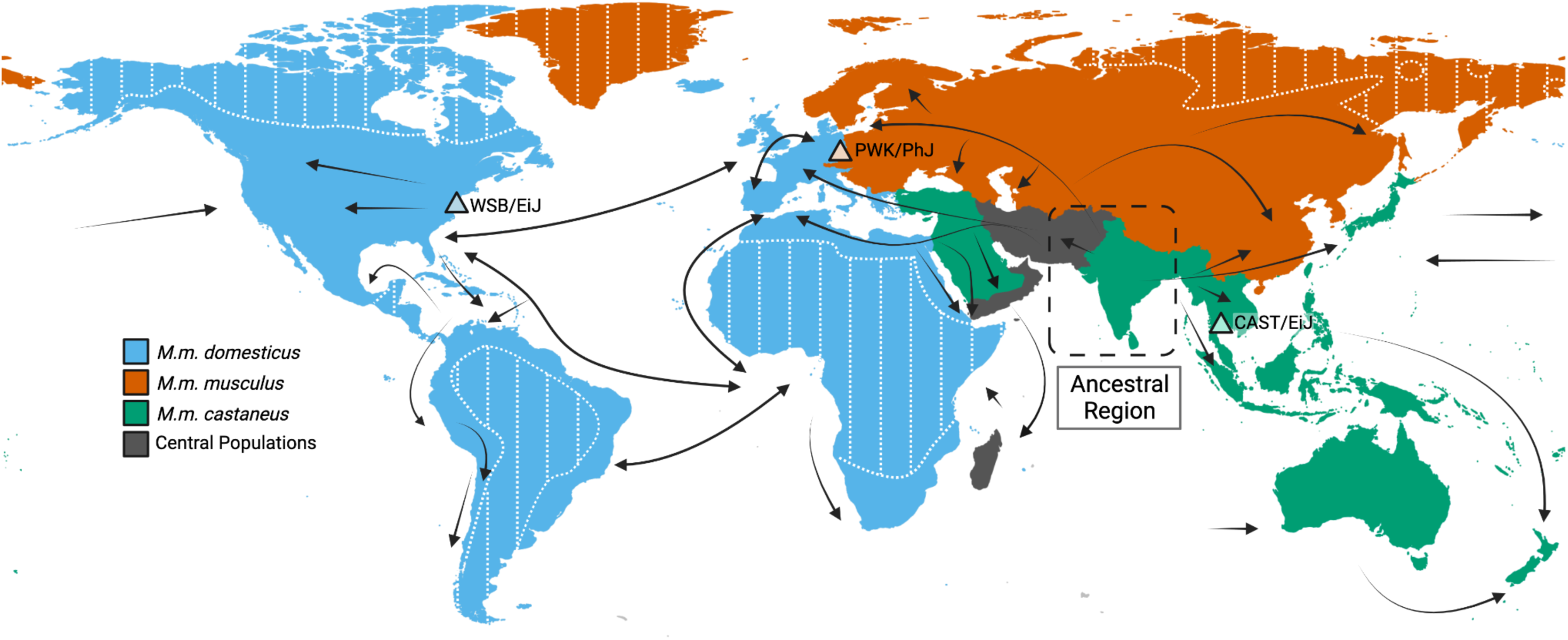
Inferred colonization history of house mice and current range map. Rough regions for each subspecies are as follows: *M.m. castaneus* is shown in green, *M.m. domesticus* is shown in blue, and *M.m. musculus* is shown in red. Importantly, this map does not include precise delineations of the boundaries of each subspecies and omits known hybrid zones. Subspecies locations may be actively in flux due to active migration and introductions. (Adapted from Guénet et al. 2012; Didion and Pardo-Manuel de Villena 2012; and Phifer-Rixey and Nachman 2015)

Human studies of phenotypes observed at the extremes of bioclimatic tolerance zones (e.g., high-altitude regions, low-precipitation regions) have been extremely fruitful in unlocking biological mechanisms of environmental resilience and adaptation [17,18]. For example, the positive selection of genes in hypoxia pathways has enabled multiple human populations to adapt to high-altitude living [18–20]. Similarly, selection for increased spleen size in breath-hold diving sea nomads [21] altered fatty acid metabolism in Arctic populations [22,23], and activation of arsenic metabolism pathways in the Atacama desert [24,25] has enabled human populations to adapt to these extreme environments. Sampling mice from their environmental extremes could likewise be a powerful strategy for discovering new, natural models of physiologically and biomedically relevant traits. In order to achieve this goal, accurate maps that delimit the geographic bounds of mouse physiological tolerance to various environmental features are needed.

The current mouse range map was developed as part of the *IUCN Red List of Threatened Species* in 2016 when *Mus musculus* was last assessed for conservation concern [26]. This range map was derived from the estimated extent of occurrence, defined as the minimum convex polygon around all present native occurrences of a species [27]. While this approach defines the general region a species may occupy, the accuracy of range estimation for species with incomplete sampling or insufficient occurrence data may be improved using alternative approaches. Further, the granularity of minimum convex polygons is limited and cannot provide insight into internal barriers to species range (e.g., mountain ranges or bodies of water that may limit dispersal) [28,29]. Lastly, there has been a significant increase in the number of occurrence records since 2016 due in large part to the increased digitization of museum collections and the availability of citizen science data [30–32]. These emergent resources have not yet been brought to bear on estimates of the house mouse species range.

Here, we present new, comprehensive geographic range maps for *Mus musculus.* We used global observations of *M. musculus* sourced from the Global Biodiversity Information Facilities (GBIF) to update the range map. Further, we perform species distribution modeling using bioclimatic variables to infer environmental tolerances and define regions where house mice may be at their physiological extremes. Across the *M. musculus* range, we predict the genetic diversity of mice based on available wild mouse genome sequencing resources. Synthesizing these outputs, we highlight under-sampled regions that may harbor high and unexplored mouse genetic diversity and areas where wild mice may be subject to unique selection pressures.

## Results

### Data Summary

A total of 108,320 *Mus musculus* occurrence records were obtained from GBIF. These records were globally distributed (Figure 2A) but concentrated in Australia, Europe, and North America (Figure 2B). The earliest records date to 1600, but most data entries were recorded within the last three decades (95% since 1991, Figure 2C). In total, 23% of records were of preserved specimens housed in museums and cultural institutions that may be suitable for DNA extraction (Figure 2D) [33]. More than half of the records, however, were human observations based on government and citizen science data sources (Figure 2D, E). Most records are not annotated to subspecies resolution (93.46%), and very few records were available for *M. m. castaneus* (n=7, Figure 2F). For these reasons, we focus on modeling only the species-level geographic range.

**Figure 2.**
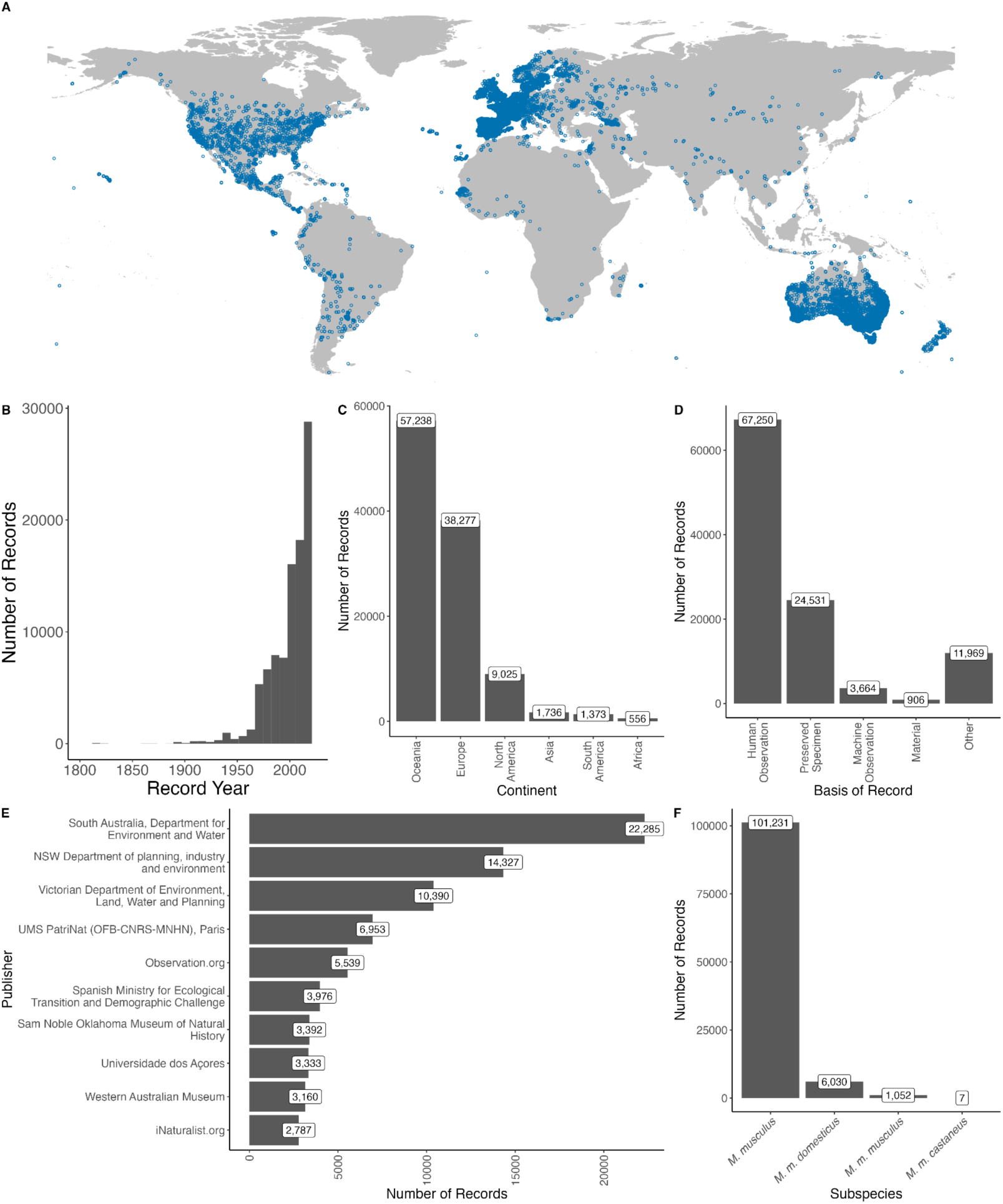
Characteristics of global house mouse occurrence records. (A) Global distribution of the 108,320 records of *Mus musculus* used in this study from the Global Biodiversity Information Facilities. (B) Distribution of the records over time clipped from 1800 to 2023. The oldest human observation was from 1600, and the oldest preserved specimen was from 1800. (C) Distribution of specimens across the continent of collection. (D) Distribution of records across the basis of record. Other bases of record include living specimens, unspecified occurrences, and unspecified observations, while ‘material’ includes both material samples and material citations. (E) Distribution of records across the top ten data providers for records. (F) Distribution of records by subspecies designation.

### Range Model

We used three range modeling methods applied to GBIF data records to update our understanding of the global distribution of house mice. The species distribution for *Mus musculus* was conducted globally and independently by continent, using the BIOCLIM algorithm, general linear modeling (GLM), and the maximum entropy (MaxEnt) algorithm. BIOCLIM is an extensively used ‘climate envelope model’ that benefits from high interpretability [34] despite generally underperforming compared to alternative approaches [35]. Unlike the other two approaches, BIOCLIM uses presence-only records and the percentile distribution of environmental variables at each presence site to find sites with predictor values close to the 50th percentile across the prediction environment. GLM is a maximum likelihood approach and is a mathematical extension of linear models. GLMs fit the mean of an outcome variable as a linear combination of predictors. GLMs have been successfully used in species distribution modeling despite being limited to linear predictor-response relationships [36]. Finally, MaxEnt is an extremely widely used model that attempts to minimize the relative entropy in covariate space between the probability density estimated from the presence data and the background landscape [37].

While all model types performed significantly better than chance (AUC > 0.5, Figure 3C), the full performance of the *M. musculus* range models varied by algorithm and continent. Across all regions, the BIOCLIM model had the lowest performance, with an average AUC of 0.794±0.057, significantly lower than the average of the GLM model (0.902±0.023, Wilcox *P*=0.001) and the MaxEnt model (0.887±0.081, Wilcox *P*=0.038). There was no significant difference (Wilcox *P*=0.53) overall between the performance of the GLM model and the MaxEnt model, but the MaxEnt model outperformed the GLM model at a global scale (0.928 vs. 0.903). When modeling each continent independently, there were no significant differences in the AUCs by continent (Kruskal-Wallis *P* = 0.25), but the lowest AUC values were seen in Asia (average AUC 0.780) and the highest for Europe (0.900).

**Figure 3.**
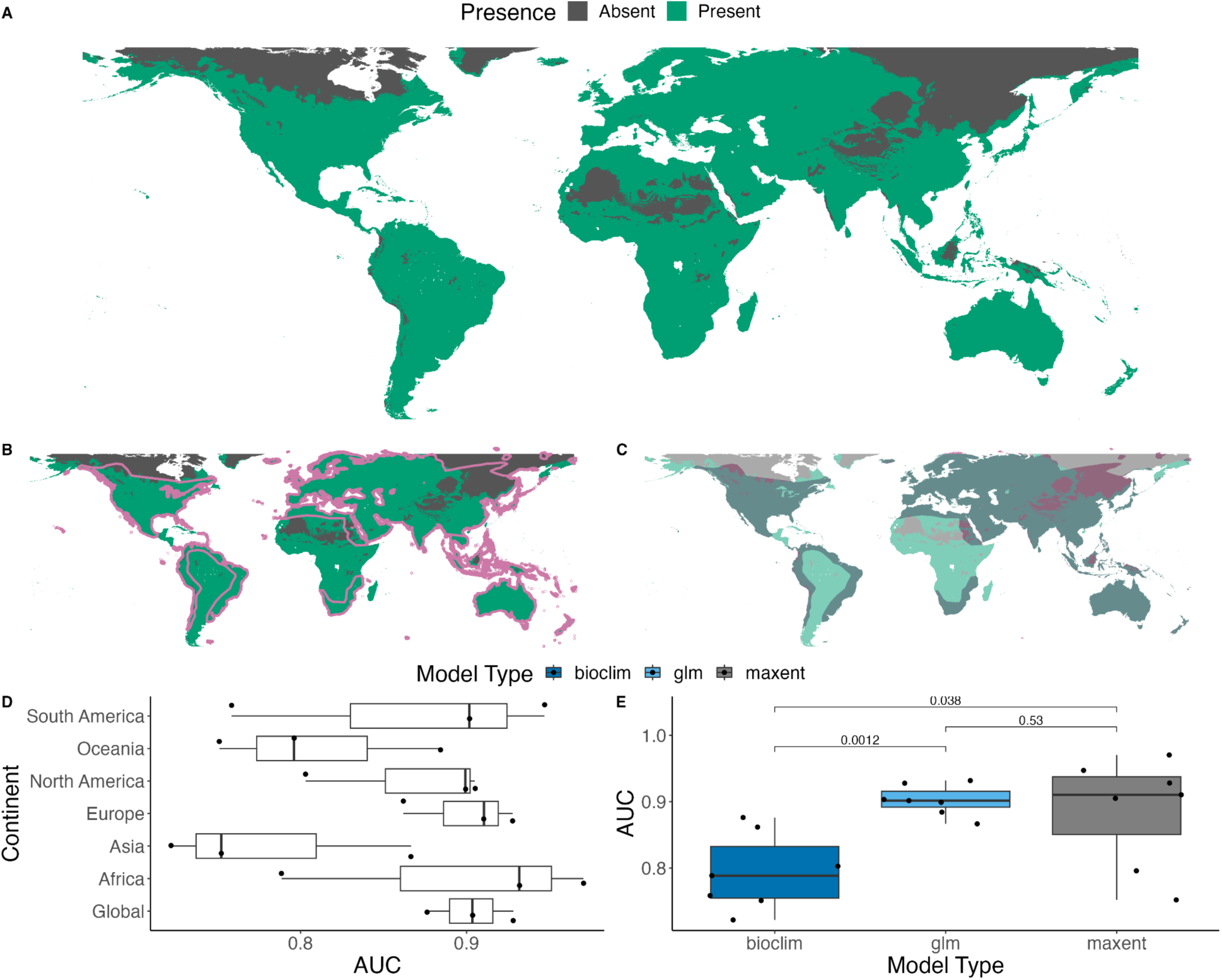
Estimated global range model for *Mus musculus*. (A) Estimated global range model based on intersecting no-omission thresholds for BIOCLIM, MaxEnt, and GLM modeling approaches. (B) A new range model overlaid with the shapefile from (Musser et al. 2021), pink. (C) Model distinctions between the previous range model and the newly presented model. Dark green areas indicate regions where the two models agree, pink indicates regions where the previous model had mouse presence, but we further delineate the range, and light green indicates new expansion areas. (D) Model performance by region, with models run independently for each region and globally. No significant differences in model performance were detected across continental regions. (E) Model performance by model type, with *P*-values reported from a Wilcox signed-rank test between model types across continents and global models.

To generate an updated range model, we intersected the three global models at a threshold that minimized the likelihood of omitting known presence points. The new *Mus musculus* range model presents several differences relative to the previous range map (Figure 3). We find a substantial increase in modeled range distribution in South America and Africa (Figure 3B-C).

North America and Europe saw only modest increases in the northern edge of their ranges. In Asia, the range area decreased, owing to further delineation of the suitable region within the previously generated maximum convex hull. Notably, the locations of this increased delineation suggest that the Himalayan mountains and the Gobi Desert may be substantial barriers to house mouse dispersal (Figure 3C). However, GBIF records in Asia were the most likely to fall in regions modeled as ‘absence’ regions in the model (6.06%, n=18 thinned omitted presence points), followed by Africa (2.70%, n=5 thinned omitted presence points). Our updated range models may be improved by increased access to occurrence data in these regions.

### Habitat Suitability and Bioclimatic Tolerances

The global *Mus musculus* model showed an extremely broad climate envelope (Figure 4A). We estimated the importance of different bioclimatic variables in defining the geographic range covered by the MaxEnt and GLM models by simulating ten independent datasets harboring random pseudo-absences (see Methods; Figure 4B).

**Figure 4.**
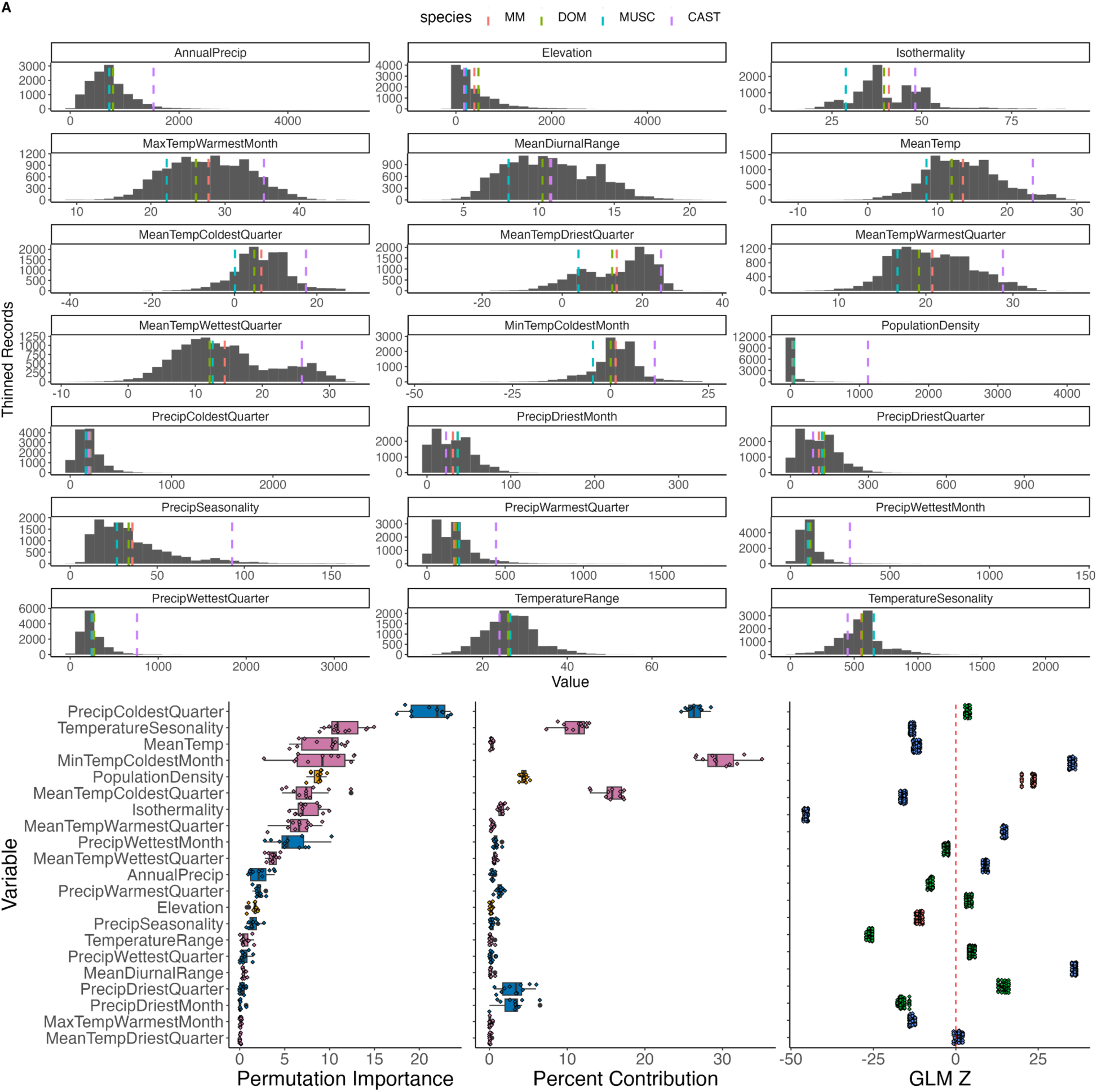
Climate variation across the newly described *Mus musculus* range. (A) Summary of species and subspecies values for presence 10-km pixels, marked with the average value across the full occurrence dataset (MM) and then independently for each subspecies (DOM, MUSC, CAST). (B) Predictor importance and contributions to the MaxEnt Model and Z-score for each predictor in the GLM model. These estimates derive from 10 permutations of the global data with re-sampled random absences and 5-fold data partitioning to generate average model values.

**Figure 5.**
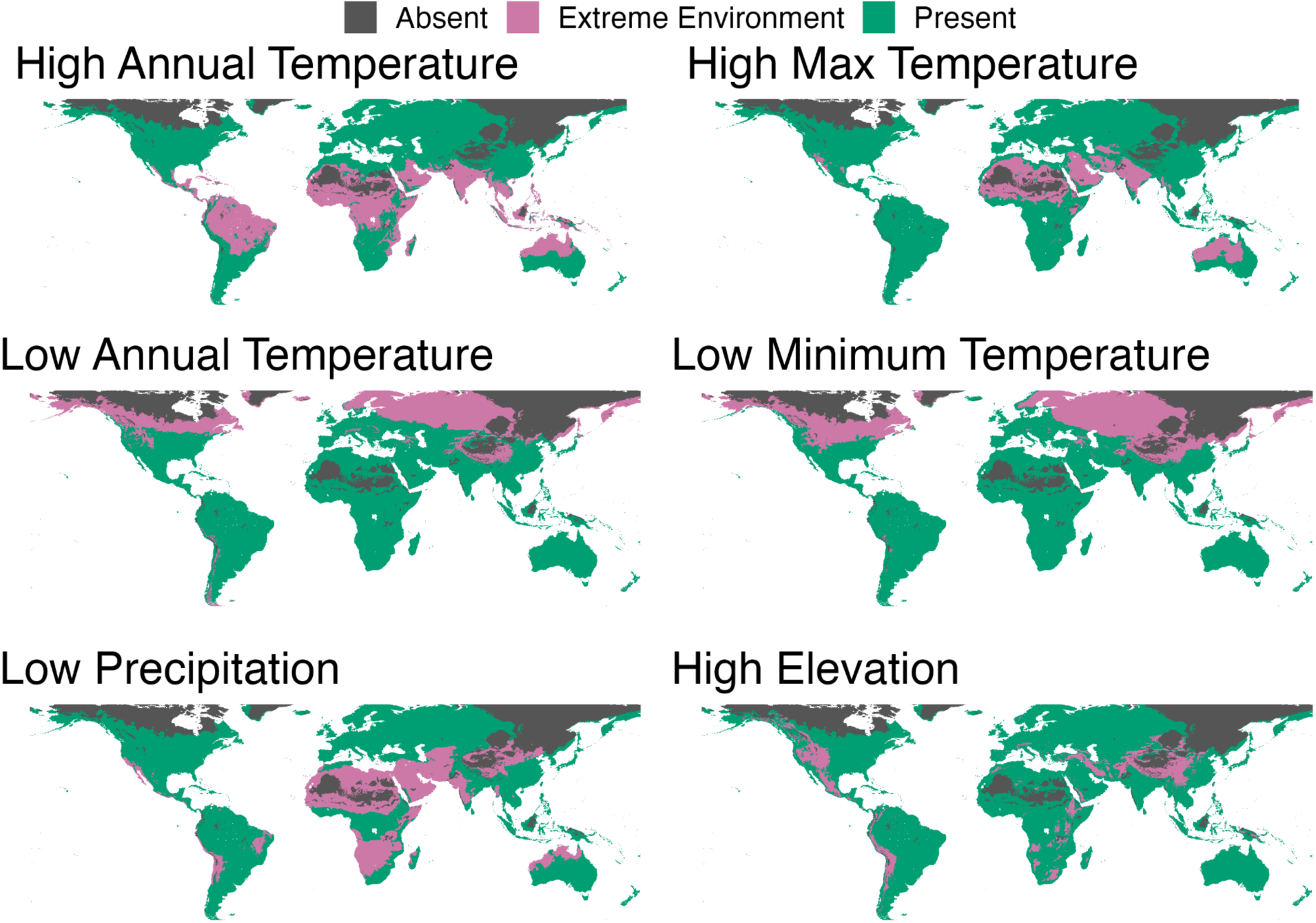
Range maps highlighting specific extremes in environmental conditions. These include high temperatures (>95th percentile of mean annual temperature and of the maximum temperature in the warmest month); low temperatures (<5th percentile of the coldest quarter and mean annual temperature); low precipitation (<5th percentile driest quarter precipitation), and high elevation (>95th percentile of elevation).

We estimated the permutation importance and percent contribution of each predictor variable in the MaxEnt model run on each simulated dataset (Figure 4B). With correlated environmental variables, the interpretation of the percent contribution from MaxEnt models can be misleading [38], so we focused on the set of predictors with the highest permutation importance. Precipitation in the coldest quarter had the largest importance and contribution to the MaxEnt models. Additional significant variables include the temperature seasonality, the annual mean temperature, and the mean temperature of the coldest month. Human population density also had a strong influence on the model, with a permutation importance of 8.67 and a percent contribution of 4.4. This finding reinforces the well-established and intimate commensal relationship between house mice and humans. We note that the correlated nature of some variables may artificially deflate their estimated influence in the MaxEnt model. For example, elevation had only a moderate permutation importance (permutation importance = 1.61, but many climatic features of high-elevation environments are captured in other predictors (e.g., mean temperature).

Similarly, we performed ten rounds of GLM training, testing, and evaluation to estimate the GLM Z-value of bioclimatic predictor variables. Increasing values of mean diurnal range (estimate = 0.107 ± 0.002, *P* < 0.0001), minimum temperature in the coldest month (estimate =0.103 ± 0.002, *P* < 0.0001), human population density (estimate = 0.001 ± 0.0001, *P* < 0.0001), precipitation in the driest quarter (estimate = 0.023 ± 0.002, *P* < 0.0001), and the mean temperature in the warmest quarter (estimate = 0.158 ± 0.005, *P* < 0.0001) were associated with a higher probability of mouse presence in the range model (Figure 4B). Alternatively, a decreasing probability of mouse occurrence was associated with increasing values of elevation (estimate = -0.0004 ± 0.00003, *P* < 0.0001), average temperature (estimate = -0.072 ± 0.005, *P* < 0.0001), temperature seasonality (estimate = -0.003 ± 0.0001, *P* < 0.0001), maximum temperature of the warmest month (estimate = -0.039 ± 0.002, *P* < 0.0001), mean temperature of the coldest quarter (estimate = -0.155 ± 0.007, *P* < 0.0001), the precipitation in the driest month (estimate = -0.072 ± 0.006, *P* < 0.0001), precipitation seasonality (estimate = -0.025 ± 0.0006, *P* < 0.0001), and isothermality (estimate = -0.194 ± 0.003, *P* < 0.0001) (Figure 4B).

Mammalian adaptation to extreme environments has been demonstrated across altitudinal, precipitation, and thermal gradients – variables that provide significant predictive power for delimiting the bounds of the mouse climate envelope. We speculate that regions at the environmental extremes of the mouse species distribution may harbor mouse populations with unique physiological, cellular, and biochemical adaptations. To identify such regions for potential future priority sampling, we generated maps highlighting regions in the house mouse species range with high temperatures (>95th percentile of mean temperature and of the maximum temperature in the warmest month), low temperatures (<5th percentile of the coldest quarter and mean annual temperature), low precipitation (<5th percentile driest quarter precipitation), and high altitude (>95th percentile of elevation) (Figure 6).

**Figure 6.**
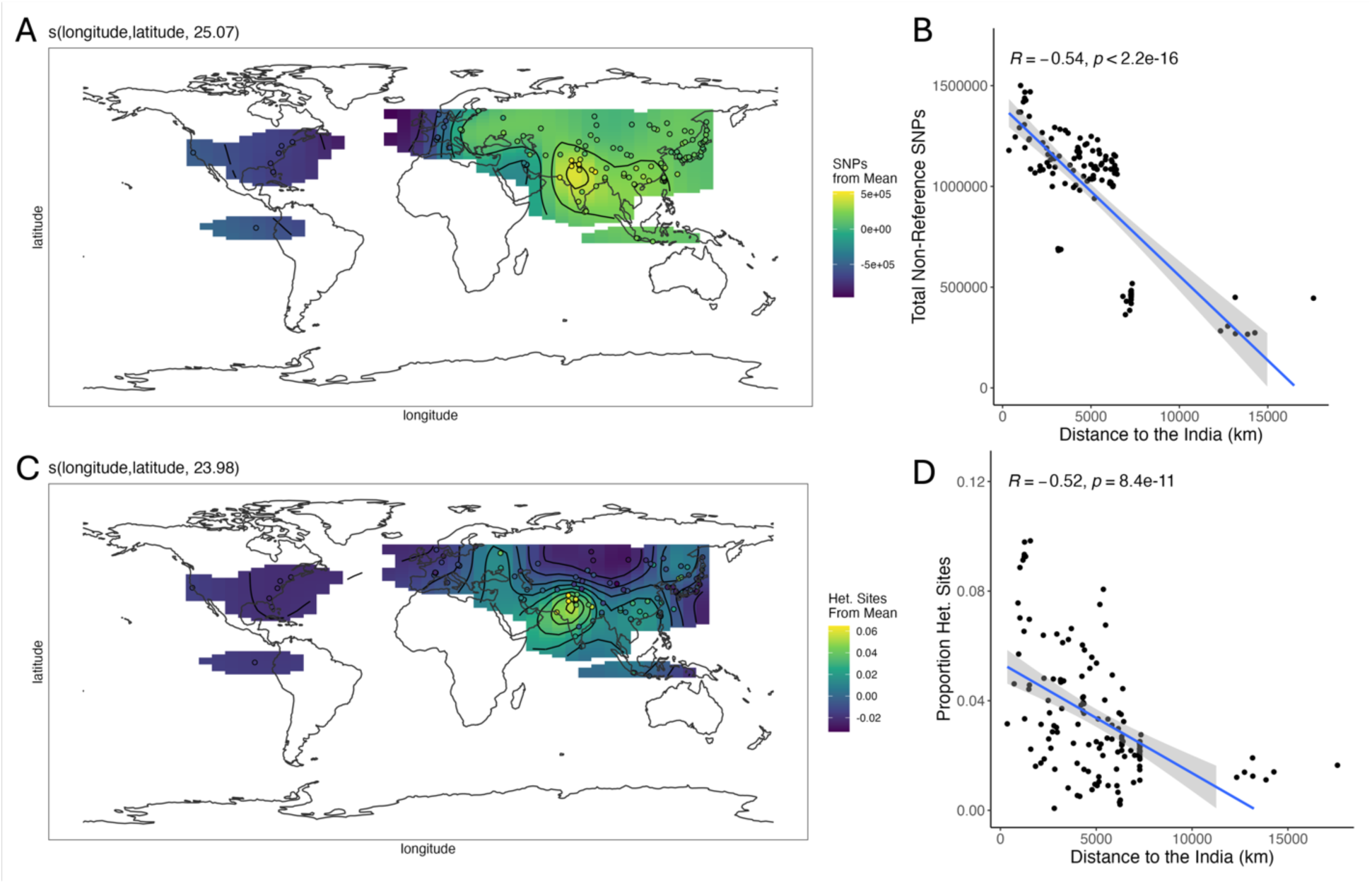
Global mouse genetic diversity. (A,C) General additive model of SNP counts plotted across the global *Mus musculus* distribution. Darker colors indicate lower than average while lighter colors indicate higher than average fitted values. Black lines demarcate contours. (B,D) Relationship between the distance to India and the total number of SNPs (B) and proportion of heterozygous sites (D), with the Pearson correlation coefficient and a fitted linear trend line.

### Predicted Genetic Diversity

Prevailing models of house mouse evolution suggest that the species emerged on the Indian subcontinent ∼0.5 MYA, although there remains scientific controversy around their precise geographic origins. Contemporary populations within a species’ ancestral region typically exhibit greater genetic diversity than populations outside the ancestral range, suggesting that maps of global genetic diversity may hold clues about the demographic history of a species.

Using publicly available whole-genome sequence data for wild-caught mice, we modeled the number of SNPs and the proportion of heterozygous sites across the *Mus musculus* range. Both metrics showed substantial variability across sequenced samples (per sample mean SNPs per chromosome: mean= 827,929± 404,127, range 1,500,598-63,920; proportion of heterozygous sites: mean=0.031±0.022, range 0.0007-0.0986).

The spatial distribution of house mice is well-captured by a general additive model, including the number of SNPs, with 94.7% of deviance explained (adjusted R^2^ = 0.936). The best-fit smoothed function of latitude and longitude had 25.07 effective degrees of freedom with an F-value of 71.27(*P* < 0.0001) and a GCV value of 8.3x10^9^. Inspection of the fitted curves suggests that the highest SNP diversity is centered in the Indian subcontinent, with decreasing diversity as samples move further into Europe (Figure 6A). Overall, the distance from India and the average total number of SNPs by chromosome are highly negatively correlated (*R*=-0.54, *P*<0.0001, Figure 6B).

An analogous model featuring the proportion of heterozygous sites as a function of spatial distribution explained 67% of the total deviance (adjusted R^2^=0.6). The best-fit smoothed function of latitude and longitude had 23.98 effective degrees of freedom with an F-value of 7.796 (*P* < 0.0001) and a GCV value of 0.0002. Peak heterozygosity localizes to the Indian subcontinent but exhibits a steeper decline when moving away from this locality (Figure 6C). As with SNP count, there was a significant negative correlation between the proportion of heterozygous sites and the distance to India (*R*=-0.52, *P* < 0.0001).

Critically, both models could not fit predictions for large portions of the *Mus musculus* range, including South America, Central America, Australia, and Africa, meaning we cannot predict genetic diversity across these regions with existing wild mouse resources (Figure 6A, C). Further, in many regions of the range where estimated house mouse density is likely to be high, model predictions of local genetic diversity are supported by a very small number of whole genome sequences. In fact, more than half of the available genomes were from Europe and Asia, with very few or no voucher samples from Africa, the Americas, or Oceania.

## Discussion

We leveraged publicly available occurrence records for house mice across the globe to update the range distribution of this important species. Our range model revealed an increase in the global distribution of house mice compared to previous maps, with especially notable expansions in South America and Africa. The range updates generated through this modeling approach may reflect changes in the distribution of house mice due to recent introductions, inland expansions, and fluctuations in population sizes, or they may simply use recently available data resources to correct inaccuracies and uncertainties in the initial mouse range model. Critically, we cannot distinguish between these possibilities and report only our best estimates of geographic ranges. Relatively coarse and climatically derived range tolerance is useful for understanding physiological thresholds and expected distributions. However, such approaches may be prone to localized errors, particularly in geographic regions where directed and targeted management activities have been particularly successful in controlling local house mouse populations or in regions where mice have direct natural competitors [39–41].

Our findings suggest that the *Mus musculus* species range is strongly delimited by environmental variables. In general, house mice are excluded from areas with extremely high temperatures, high elevation, low temperatures, or low precipitation. Many of these specific environmental limits are also limits to human tolerance [17,24,42] and, therefore, may reflect their commensal relationship with humans [43,44]. Indeed, we found an increasing likelihood of mouse occurrence associated with increasing human density. At the same time, our work also spotlights some extreme environments where house mice and humans coincide and where parallel adaptation in both species may be mediated by genetic changes in the same genes or pathways [45].

We also show that the *Mus musculus* species distribution is well-predicted by diurnal range and isothermality, metrics that quantify daily and annual temperature fluctuations, respectively. Previous work suggests that habitat generalists like *Mus musculus* tend to thrive under high temperature variability [46]. Consistent with these earlier findings, increasing diurnal range and decreasing isothermality (*e.g.,* more variable daily and seasonal temperatures) were strongly associated with a higher probability of mouse occurrence (Figure 4B). Anthropogenic climate change is projected to increase seasonal variability [47]. Thus, understanding the capacity for adaptation to changing isothermality and diurnal temperature changes will likely provide substantial insight into how organisms adapt to climate change. Indeed, recent work in humans has suggested that diurnal temperature range shifts will be an increasingly significant driver of human climate-related mortality [48–51].

We utilized existing WGS for wild-caught mice to estimate wild mouse genetic diversity in our updated species range. We observe the highest SNP diversity and heterozygosity centered on the Indian subcontinent (Figure 6), supporting the hypothesis that *Mus musculus* originated in this region [52,53]. The significant negative correlation between the number of SNPs and the proportion of heterozygous sites with distance from India further strengthens this assertion. However, we currently lack sufficient whole genome sequence resources for samples in large portions of the *Mus musculus* range, including South America, Central America, Australia, and Africa, limiting our ability to estimate Mus musculus genetic diversity in these regions reliably. These findings underscore the need for more comprehensive sampling and sequencing efforts across the global range of *Mus musculus*. Such efforts would not only enhance our understanding of mouse genetic diversity but also provide valuable insights into house mouse demographic history and adaptation.

One limitation of the range modeling approach employed here is the reliance on occurrence records and existing literature, which introduces biases and incomplete data. In particular, our range map boundaries are imprecise due to under-sampling at environmental extremes. Further, the assignment of mouse records to the subspecies level is rare, which precluded the development of subspecies-specific models. Together, these limitations emphasize the importance of continued field sampling of wild mice and ecological studies at the periphery of the mouse range. Government sampling programs may provide especially powerful sources of additional structured sampling. Indeed, nearly 50% of the observations included in our study are provided to GBIF by government programs. Many of these data programs collect additional information during the data collection projects, including temporal presence/absence data and various phenotyping data, including the species, age, sex, weight, and body condition of the organism. Additionally, data may include information on habitat characteristics, such as vegetation type and structure, which can be important for understanding the ecological context of house mouse occurrences. Including these phenotypes and structured sampling information alongside occurrences will improve our understanding of how the phenotype and ecological context of mice vary across their global range. Citizen monitoring and community science projects have also made substantial contributions to house mouse occurrence data records. Approximately 10% of the observational data used here are from two large citizen science projects, Observation.org and iNaturalist (Figure 2E). Because humans often encounter house mice across their range, encouraging documentation of these interactions through these community platforms will provide needed data to further refine species range estimates and inform on the frequencies of human-mouse interactions.

In addition to providing a foundation for future ecological studies, estimating the global distribution of house mice and their environmental envelope can guide new sampling efforts aiming to uncover unique genetic and functional diversity. House mice are an important biomedical model system, but laboratory mice sample a modest proportion of the genetic and phenotypic diversity observed in the wild. This consideration inherently restricts the scope of possible biological discovery in laboratory mouse models and limits efforts to bridge the mouse-human translational interface. Our work not only illuminates the geographic regions with the highest levels of diversity but also spotlights regions at the extremes of the environmental envelope where genetic adaptation has potentially led to novel phenotypes. Local adaptive evolution is a major driver of phenotypic divergence between human populations, including traits associated with disease susceptibility and resilience [54]. Thus, our range maps may help guide the development of naturalistic mouse models that more accurately reflect the evolutionary origins of human disease, complementing conventional methods of mouse model generation reliant on targeted genetic engineering of inbred strains.

## Conclusion

We provide comprehensive range maps and species distribution models for *Mus musculus.* The updated range maps yield substantial increases in the range area of house mice compared to previous range maps, particularly in South America and Africa. Additionally, our work offers predictions for the level of genetic diversity across the mouse range, with the highest diversity observed in the Indian subcontinent, which aligns with prior work defining this region as the ancestral epicenter of *Mus musculus*. Regions of predicted high genetic diversity, as well as regions of recent expansion or at the limits of bioclimatic tolerances, are underrepresented in existing wild mouse genome sequencing data but may harbor unique genetic and functional diversity. Overall, our work refines the global house mouse species range, defines parameters of habitat suitability, and estimates the level of genetic diversity in house mouse populations across the globe. Collectively, these new insights stand to inform future studies of wild mouse ecology and population genetics.

## Methods

### Species occurrence records

Occurrence records for *Mus musculus*, *M. m. domesticus*, *M. m. castaneus*, and *M. m. musculus* were downloaded from the Global Biodiversity Information Facility (GBIF: https://www.gbif.org/occurrence/download/0051955-231002084531237, Figure 2). The collection timeframe for GBIF records extended from 1800 to 2023, with 50% falling between 1989 and 2016. A total of 108,320 occurrences were available, with coordinates and uncertainty under 100. After thinning to a 10-km radius using spThin in R [55] and removing duplicates, a total of 13,126 *Mus musculus* records were available, including 6 *M.m. castaneus*, 1,188 *M.m. domesticus* records, and 199 *M.m. musculus* records. The low number of records for the three primary subspecies precluded modeling at the subspecies level.

### Bioclimatic and predictor variables

BIOCLIM variables were downloaded using the dismo package in R from CRU-TS 4.06 [56] and downscaled with WorldClim 2.1 [57]. These include 22 environmental covariates at 2.5 arc-second resolution, including precipitation, temperature, and seasonality (Table 1). We also used the gridded global bedrock elevation model to explicitly model high-elevation effects that may not be fully captured by the climate predictors [58]. Because house mice are commensal with human populations, we also included the human population density as an additional predictor variable [59].

**Table 1.** Summary of predictor variables. Distribution of occurrence records across climate variables, elevation, and population density. Occurrence records thinned to 1 per 10 km cell prior to analysis (n=13,126). A quarter is ¼ of the year or three months. All numbers are presented as Mean ± Standard Deviation (5% - 95%).

| Variable | Mean ± Standard Deviation (5% - 95%) |
| --- | --- |
| **Temperature** |  |
| Mean Temperature | 136.43 ± 51.9 (58 - 228) °C * 10 |
| Mean Diurnal Range<br>*Mean of monthly (max temp - min temp)* | 107.29 ± 28.49 (64 - 156) °C * 10 |
| Isothermality<br>*(Mean Diurnal Range / Temperature Range) × 100* | 40.84 ± 9.77 (26 - 55) |
| Temperature Seasonality<br>*standard deviation × 100* | 5566.22 ± 1818.04 (2851.7 - 8887.05) °C * 10 |
| Max Temperature (Warmest Month) | 277.58 ± 54.83 (193 - 367) °C * 10 |
| Minimum Temperature (Coldest Month) | 12.84 ± 61.15 (-97 - 100) °C * 10 |
| Temperature Range | 264.75 ± 62.46 (169 - 375) °C * 10 |
| Mean Temperature (Wettest Quarter) | 143.89 ± 72 (44 - 277.05) °C * 10 |
| Mean Temperature (Driest Quarter) | 136.37 ± 87.72 (-22 - 246) °C * 10 |
| Mean Temperature (Warmest Quarter) | 207.42 ± 45.34 (140 - 287) °C * 10 |
| Mean Temperature (Coldest Quarter) | 65.2 ± 65.23 (-45 - 167) °C * 10 |
| **Precipitation** |  |
| Annual Precipitation | 720.15 ± 391.71 (218 - 1386) mm |
| Precipitation (Wettest Month) | 95.16 ± 61.07 (30 - 197) mm |
| Precipitation (Driest Month) | 31.11 ± 23.39 (2 - 72) mm |
| Precipitation Seasonality<br>*Coefficient of Variation* | 35.75 ± 23.04 (12 - 85) mm |
| Precipitation (Wettest Quarter) | 258.51 ± 162.75 (78 - 536) mm |
| Precipitation (Driest Quarter) | 109.7 ± 75.63 (13 - 242) mm |
| Precipitation (Warmest Quarter) | 174.37 ± 131.8 (28 - 413) mm |
| Precip (Coldest Quarter) | 179.66 ± 121.32 (32 - 387) mm |
| **Geographic** |  |
| Population Density | 311.8 ± 1270.2 (0 - 1545.17) |
| Elevation | 380.56 ± 472.18 (1.08 - 1316.1) m |

### Species distribution modeling

The species distribution for *Mus musculus* was conducted globally and independently by continent, using the BIOCLIM algorithm, general linear modeling (GLM), and the maximum entropy (MaxEnt) algorithm. Model fitting and analysis was performed using the dismo R package [60] and visualized using sf [61,62], ggpubr [63], maptools [64], and the Tidyverse suite [65].

Because the occurrence data represented presence-only data (i.e., no true absence data is available), pseudo-absences are needed to differentiate between localities where house mice are known to occur and regions where no presence points have been observed. Therefore, alongside the spatially-thinned presence records for *Mus musculus*, pseudo-absences were generated randomly from the background. The number of random pseudo-absences should be approximately equal to three times the number of presence points used in the model [66]. Guided by this threshold, we generated pseudo-absences independently for each continent and for the global models. GLM and MaxEnt models were permuted ten times to evaluate predictor stability, with random pseudo-absences independently generated for each permutation.

The performance of each model was evaluated using both internal and external test data randomly partitioned from presences and pseudo-absences. We used AUC (Area Under the Receiver-Operator Curve), measured with the internal and external test data to determine the model performance. An AUC of 0.5 indicates the model is no better than chance, but high values of AUC indicate that regions with high predicted suitability tend to be regions with known presence, while regions with lower suitability predictions tend to be regions where pseudo-absences and few presence records fall. To generate an average model, the predictions for the GLM, BIOCLIM, and MaxEnt models were normalized to fall between 0 and 1 and weighted by their AUC scores. To generate the weights, we use the following formula to assign higher weights to higher AUC values: *weight* = (*AUC* − 0.1)^2^. To generate the final range map, each model’s highest threshold at which there was no omission of training presence points was intersected to provide a conservative estimate of mouse absence regions.

### Predicting Global Mouse Nucleotide Diversity

A total of 267 wild mouse samples were used to investigate genetic diversity across the mouse range. Sample accession numbers for the wild mouse genome sequences used are provided in Supplemental Table 1. Reads were mapped to the GRCm39 and DeepVariant (v1.2.0 [67]) using “WGS” mode. The resulting gVCF files were then merged using glnexus (v1.2.7 [68]) using the DeepVariantWGS configuration to produce per-chromosome joint call sets. Sites with >10% missing data, genotype quality <30, and indels were subsequently filtered using bcftools (v 0.1.19; [69]). The bcftools stats command was used to obtain per-sample counts of SNPs for each chromosome. The proportion of heterozygous sites was calculated as the ratio of heterozygous sites to all successfully genotyped sites, and the total number of SNPs was calculated as the sum of the heterozygous and non-reference homozygous sites. For locations with whole-genome sequence data for more than one wild mouse, SNP counts and heterozygosity estimates were averaged across samples. Data were available for a total of 138 unique locations. To model the distribution of the number of SNPs and the proportion of heterozygous variants across these spatial points, we used a general additive model with a smoothed joint predictor of latitude and longitude fit to the genetic diversity metrics in R (v4.4.1) using mgcv (v1.9-1, [70–72]) and mgcViz (v0.1.11, [73]). The model was fit as follows:

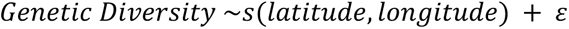

The resulting spatial distribution was used to visualize regions where genetic diversity may be highest. Additionally, we computed the geographic distance between each location and the center point of India (20.59 N 78.96 E) on the WGS ellipsoid using the pointDistance function from the raster package in R (v3.6-26, [74]). Pearson’s correlation was used to assess the relationship between SNP count and heterozygosity and the distance from India.

---

Supplemental Table 1. Accession numbers and genome diversity values for wild mouse samples.

